# Intercellular communication in the fern endosymbiotic cyanobacterium *Nostoc azollae*

**DOI:** 10.1101/2025.04.17.649285

**Authors:** Cristina Sarasa-Buisán, Mercedes Nieves-Morión, Peter Lindblad, Sandra Nierzwicki-Bauer, Henriette Schlüpmann, Enrique Flores

## Abstract

The water fern *Azolla* spp. harbors as an endobiont the N_2_-fixing, filamentous, heterocyst-forming cyanobacterium *Nostoc azollae. N. azollae* provide the fern with fixed nitrogen permitting its growth in nitrogen-poor environments. In the filaments of heterocyst-forming cyanobacteria, an intercellular exchange of regulators and metabolites occur in which heterocysts provide vegetative cells with fixed nitrogen and vegetative cells provide heterocysts with reduced carbon. Intercellular molecular exchange takes place by diffusion through septal junctions and can be probed by fluorescence recovery after photobleaching (FRAP) analysis with fluorescent markers such as calcein and 5-carboxyfluorescein. The septal junctions traverse the septal peptidoglycan through nanopores that can be visualized in isolated septal peptidoglycan disks by electron microscopy. Here we obtained *N. azollae* material from *Azolla* plants, which contains the symbiotic cyanobacterium in a viable state and with different morphologies, including heterocyst-containing filaments. FRAP analysis showed effective transfer of the fluorescent markers between vegetative cells as well as from vegetative cells to heterocysts. Interestingly, communicating and noncommunicating vegetative cells and heterocysts could be distinguished showing conservation in the endobiont of a regulatory mechanism capable of opening and closing septal junctions. Peptidoglycan sacculi were also isolated and showed septal disks with arrays of nanopores that conform to those visualized in other heterocyst-forming cyanobacteria. However, a wider range of septal disk size was observed in *N. azollae*. In spite of its eroded genome, *N. azollae* maintains the intercellular communication system that is key for its growth as a multicellular organism.

**Importance:** The water fern *Azolla* constitutes a unique symbiotic system in which cyanobacterial endobionts capable of fixation of atmospheric nitrogen provide the plant with the nitrogen needed for growth. This symbiosis is an important fertilizer for rice crops worldwide, thereby reducing the reliance on fossil fuel-derived nitrogen fertilizers. The symbiotic cyanobacterium, *Nostoc azollae*, is a heterocyst-forming strain in which a filament of cells is the organismic unit of growth. Here we show that the intercellular molecular exchange function necessary for the multicellular behavior of the organism is conserved in the endobiotic *Nostoc azollae*.

## Introduction

The water fern *Azolla* spp. is characterized by harboring as an endobiont the N_2_-fixing, filamentous, heterocyst-forming cyanobacterium *Nostoc azollae* (1, 2). The heterocysts are cells specialized for the fixation of atmospheric N_2_, and the endobiont allows the growth of the fern in the absence of any source of combined nitrogen (3). Cyanobacteria are characterized by performing oxygenic photosynthesis, and the fern additionally contains heterotrophic bacteria whose function in the symbiosis remains to be fully elucidated (4). *N. azollae* is transmitted vertically and contains an eroded genome of about 5.49 Mbp (5). *N. azollae* within the fern exhibits extensive morphological diversity, including vegetative filaments (with or without heterocysts), hormogonia (which lack heterocysts), and akinetes (spore-like cells). The developmental cycles of the fern and its endobiont are tightly synchronized. *N. azollae* originates from the apical meristems of *Azolla* in which it forms the *Shoot Apical Nostoc Colony* (SANC), where small-celled motile filaments without heterocysts (“hormogonia”) seem to be initially present (6). As the fronds grow, *N. azollae* colonizes the leaf pocket of developing leaves that emerge just behind the meristematic tip, where *N. azollae* starts to form heterocysts. The frequency of heterocysts increase along with the leaf age, reaching frequencies of 20% to 30% in filaments, which is higher than that found in free-living *Nostoc* or *Anabaena* spp. (7, 8). Moreover, behind the meristematic tip, the megasporocarp is formed and colonized by symbiotic filaments. In megasporocarps, endobiont cells differentiate into akinetes that later germinate as new *Azolla* sporelings emerge, colonizing newly developed apical meristems (9).

Fern-associated nitrogenase activity has been reported (3) and, actually, leaf nitrogen fixation rates respond to a developmental gradient, where mature leaves have higher heterocyst frequencies and N_2_ fixation rates (10, 11). However, after the 20^th^ leaf of the frond, the heterocysts begin to senesce and no longer show increased N_2_ fixation (7, 12). The nitrogenase enzyme component Fe-protein (NifH) has been detected by immunogold localization in the endobiont (13). Additionally, cyanobacterial filaments isolated from the fern have been used to study transcript levels of key genes related to carbon and nitrogen assimilation, and transcripts of *nifHDK* encoding nitrogenase were indeed detected (14). Isolated filaments were also used in studies on [^13^N]N_2_ fixation showing that *N. azollae* (then called *Anabaena*) accumulates more ammonium ions than free-living cyanobacteria, suggesting that ammonium is the N-containing compound transferred from the endobiont to the fern (15).

The growth of *N. azollae* with heterocyst-containing filaments, would suggest that intercellular molecular exchange (also called intercellular communication) is needed along the filament, as is the case for free-living heterocyst-forming cyanobacteria (16). Intercellular communication in filamentous cyanobacteria can be studied by Fluorescence Recovery After Photobleaching (FRAP) analysis (17). Two commonly used fluorescent tracers are calcein (622.5 Da) and 5-carboxyfluorescein (5-CF; 376.3 Da), both of which are negatively charged (24). Intercellular transfer takes place by simple diffusion (18) through proteinaceous structures known as septal junctions (19, 20). Cyanobacteria are diderm bacteria, and in filamentous cyanobacteria the outer membrane does not enter the septum between cells, whereas each cell is surrounded by its cytoplasmic membrane and peptidoglycan (PG) (21). Septal junctions traverse the septal PG by holes termed nanopores that can be readily seen in isolated murein (PG) sacculi (22, 23)

Intercellular molecular exchange is necessary for mutual nutrition between CO_2_-fixing vegetative cells and N_2_-fixing heterocysts to allow growth, and it may also be necessary for an efficient transfer of nitrogen to the host fern. Here, we show effective intercellular exchange of fluorescent markers and the presence of septal PG nanopores in filaments of *N. azollae* freshly isolated from *Azolla filiculoides*.

## Results and Discussion

### *Azolla* juice contains metabolically active *Nostoc azollae* cells

Material from *Azolla filiculoides* referred to as “*Azolla* juice” was isolated as described in Materials and Methods (see also Suppl. Fig. S1) and found to be enriched in *Nostoc azollae* (Fig. 1). The wide variety of *N. azollae* cells and filaments observed likely reflects the fact that, using this extraction method, the isolated cyanobiont originates from all the different tissues of the plant. Therefore, we observed *N. azollae* cells in different developmental stages, i.e., they can be akinetes, filaments of vegetative cells with or without heterocysts, or “hormogonia”-like filaments with narrowed cells (9). Additionally, the juice contained bacteria which are part of the symbiosis (24)and eukaryotic algae that may be present on the surface of the fern.

**Figure 1.**
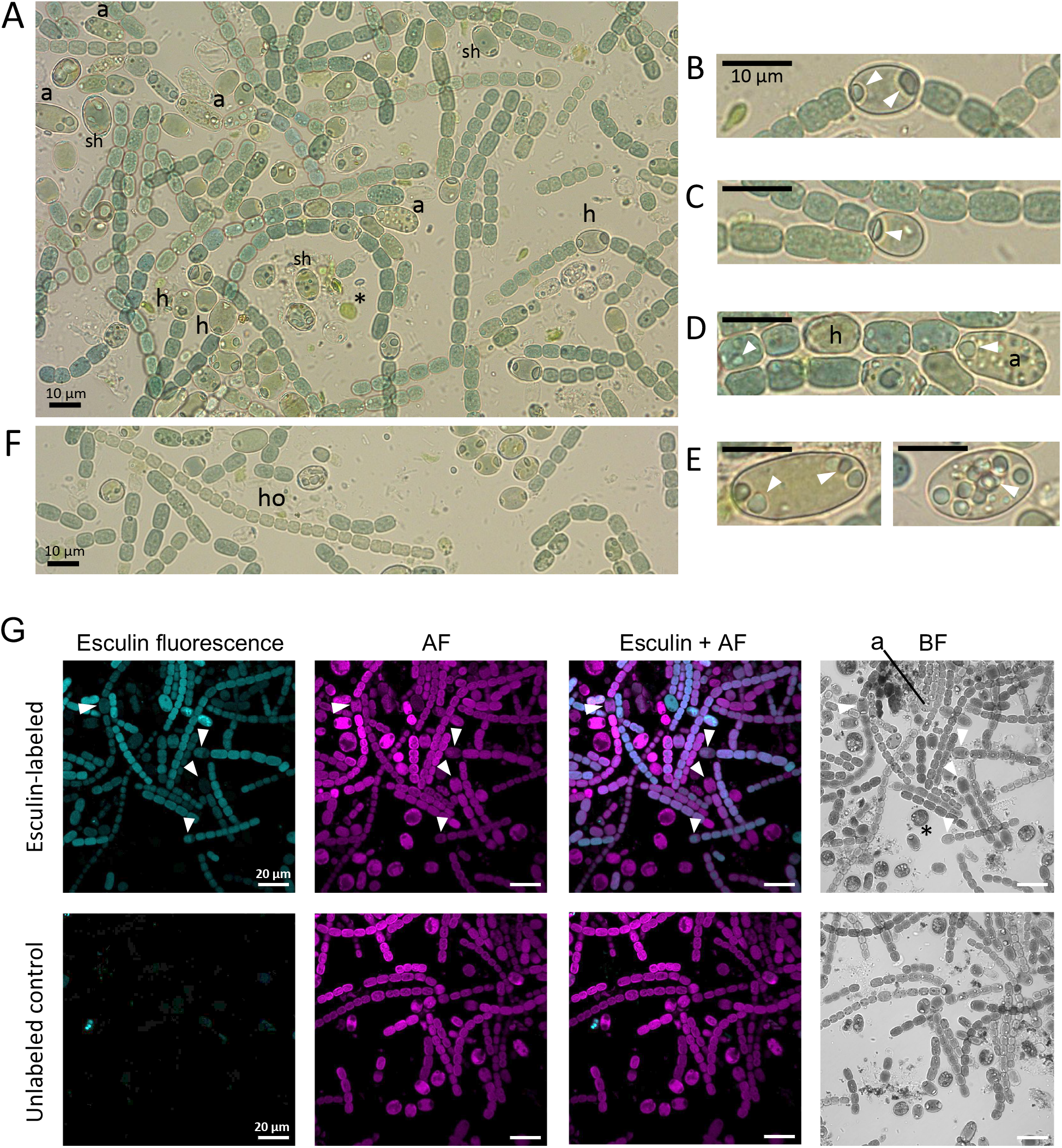
*Nostoc azollae*-enriched *Azolla* juice contains metabolically active *N. azollae* cells. (A) Brightfield micrograph of recently isolated *Nostoc azollae* from *Azolla* juice depicting the variability in morphology and cell differentiation of the cyanobacterium, with presence of heterocysts (h) and akinetes (a) at various stages of formation. Cyanophycin granules are prominent, especially at the poles of heterocysts. Note the presence of single heterocysts (sh), which are likely derived from mechanical fragmentation of filaments. Eukaryotic algae (*), which were likely present on the surface of the fern, and bacteria are also present. (B-E) Close-up of heterocysts and akinetes showing cyanophycin granules (some indicated by white arrowheads): (B) intermediate heterocyst; (C) terminal heterocyst; (D) filament with an intermediate heterocyst (h) and an akinete in formation (a), with granules visible in vegetative cells; (E) two akinetes, one with polar granules (left) and one with granules distributed throughout (right). (F) Example of a hormogonium-like filament (ho), with narrowed cells and no heterocysts. (G) Uptake of esculin by *N. azollae* filaments in leaf juice. (*Top row*) Esculin-labeled cells, with some heterocysts indicated by white arrowheads; a, akinete. (*Bottom row*) Control with unlabeled cells. Esculin fluorescence was induced at 335 nm and acquired between 443 and 490 nm; AF, autofluorescence; BF, bright field. Scale bars for (A-F) are 10 μm and for (G), 20 μm.

To study the metabolic state of the *N. azollae* cells in the juice, the filaments were incubated with esculin, a fluorescent sucrose analog that is actively taken up into the cyanobacterial cells by ABC glucoside transporters (25– 27). Vegetative cells, heterocysts and some akinetes were labeled well with esculin (Fig. 1G). Since ABC transporters require ATP, this indicates that the cells were active. In contrast, the eukaryotic algae observed in the samples were not labeled, likely reflecting lack of appropriate membrane transporters. BlastP analyses revealed that the genome of *N. azollae* encodes orthologs of some of the components of ABC glucoside transporters known to move esculin into the cells of the model heterocyst-forming cyanobacterium *Anabaena* (26, 28): GlsC and GlsD (nucleotide binding domain [NBD] proteins), and GlsP and GlsQ (transmembrane domain [TMD] proteins) (Table 1). The *N. azollae* genome also encodes a periplasmic substrate-binding protein (SBP), similar to All1027 of *Anabaena* (24), that is homologous to that of ABC disaccharide transporters. Moreover, from the two sucrose-splitting invertases described to be involved in sucrose catabolism in cyanobacteria (29, 30), *N. azollae* conserves one that is most similar to InvB, which is known to be expressed mainly in heterocysts (29, 30) (Table 1). Labeling with the sucrose analogue esculin and the presence of a putative sucrose transporter and of an invertase suggest that *N. azollae* is supplied with sucrose by the host fern. This must be important to support N_2_ fixation in filaments that contain a high percentage of heterocysts, i.e., a low percentage of CO_2_-fixing vegetative cells. Indeed, sucrose is the main photosynthetic product in *Azolla* (31).

**Table 1.** Identification of potential orthologues in the *N. azollae* genome for proteins involved in septal junctions, nanopore array machinery, glucoside ABC transporters, and sucrose metabolism. Analyses were performed using BlastP with default parameters (% query coverage and % identity shown) with FASTA sequences of the indicated protein from *Anabaena* sp. PCC 7120 as the query and the *N. azollae* genome annotation from March 28, 2024. (*) No potential orthologue found. (**) In case of InvA, the only similar protein encoded in the *N. azollae* genome is InvB, and in case of Alr2722, the only similar protein encoded in the *N. azollae* genome is the homologue of All1027 (SBP). A major facilitator superfamily protein, HepP, involved in glucoside uptake in *Anabaena* (26) has no clear homologue in *N. azollae*.

| Protein name | ORF PCC 7120 | Length (aa) | Reference | Potential orthologue in *N. azollae* | Length (aa) | Query Cover | Identity |
| --- | --- | --- | --- | --- | --- | --- | --- |
| **Proteins described to be involved in septal junction and nanopore array formation** |  |  |  |  |  |  |  |
| SepJ | *alr2338* | 751 | (32) | AAZO_RS17720 | 728 | 99% | 50.33% |
| FraC | *alr2392* | 179 | (22, 33) | AAZO_RS00195 | 182 | 99% | 61.24% |
| FraD | *alr2393* | 343 | (22, 33) | AAZO_RS00190 | 336 | 99% | 62.65% |
| FraE | *alr2394* | 267 | (22, 33) | AAZO_RS00185 | 259 | 97% | 68.73% |
| FraI | *alr4714* | 232 | (47) | AAZO_RS10910 | 210 | 94% | 60.55% |
| HglK | *all0813* | 727 | (48) | AAZO_RS07385 | 706 | 100% | 62.02% |
| SjcF1 | *all1861* | 269 | (35) | AAZO_RS24465 | 197 | 94% | 51.78% |
| SjdR | *alr0248* | 218 | (36) | * |  |  |  |
| SepI | *alr3364* | 521 | (49) | AAZO_RS22910 | 497 | 89% | 50.41% |
| SepT | *all2460* | 426 | (50) | AAZO_RS06650 | 420 | 99% | 55.79% |
| SepN | *all4109* | 235 | (34) | AAZO_RS15255 | 238 | 99% | 56.49% |
| AmiC1 | *alr0092* | 627 | (43) | AAZO_RS04180 | 619 | 100% | 64.42% |
| AmiC2 | *alr0093* | 627 | (43) | AAZO_RS04175 | 631 | 100% | 67.87% |
| LytM factor | *alr3353* | 760 | (42) | AAZO_RS21965 | 788 | 99% | 50.06% |
| FurC | *alr0957* | 149 | (51) | AAZO_RS01485 | 145 | 96% | 82.52% |
| **Glucoside ABC transporter proteins and sucrose metabolism** |  |  |  |  |  |  |  |
| GlsC (NBD) | *alr4781* | 432 | (26) | AAZO_RS14955 | 427 | 99% | 77.47% |
| GlsP (NBD) | *all0261* | 276 | (26) | AAZO_RS16245 | 277 | 96% | 37.74% |
| GlsD (TMD) | *all1823* | 366 | (28) | AAZO_RS14955 | 427 | 99% | 42.82% |
| GlsQ (TMD) | *alr2532* | 301 | (28) | AAZO_RS23735 | 310 | 93% | 35.42% |
| GlsR (SBP) | *all1916* | 418 | (28) | * |  |  |  |
| All1027 (SBP) | *all1027* | 432 | (28) | AAZO_RS01530 |  | 100% | 82.18% |
| Alr2722 (SBP) | *alr2722* | 432 | (28) | ** |  | 94% | 22.20% |
| Alr4277 (SBP) | *alr4277* | 461 | (28) | * |  |  |  |
| InvA | *alr1521* | 468 | (29) | ** |  |  |  |
| InvB | *alr0819* | 483 | (29) | AAZO_RS22440 | 482 | 98% | 83.40% |

### Freshly isolated *N. azollae* show intercellular communication

BlastP analysis showed that *N. azollae* conserves a number of the key proteins involved in intercellular molecular transfer described in *Anabaena*, such as SepJ (32), FraCDE (22, 33), SepN (34) and SjcF1 (35) among others, although it does not conserve the SjdR protein involved in the structural definition of the intercellular septa (36) (Table 1). To test the potential performance of *N. azollae* in intercellular molecular transfer, FRAP analysis was set up with the cyanobacterial filaments in *Azolla* juice.

Calcein AM and 5-carboxyfluorescein AM are hydrophobic compounds that enter the cells by diffusion through the cytoplasmic membrane. In living cells, these compounds are hydrolyzed by cytoplasmic esterases, so they can also be used as probes for discriminating live or dead cells (37). The products of the hydrolysis, calcein and 5-CF, are hydrophilic and highly fluorescent. Cyanobacterial filaments in the *Azolla* juice were well labeled with calcein and 5-CF (see “Pre” in Fig. 2A and 3A), consistent with cells being alive corroborating the results with esculin described above. This provided filaments that were suitable for FRAP analysis.

**Figure 2.**
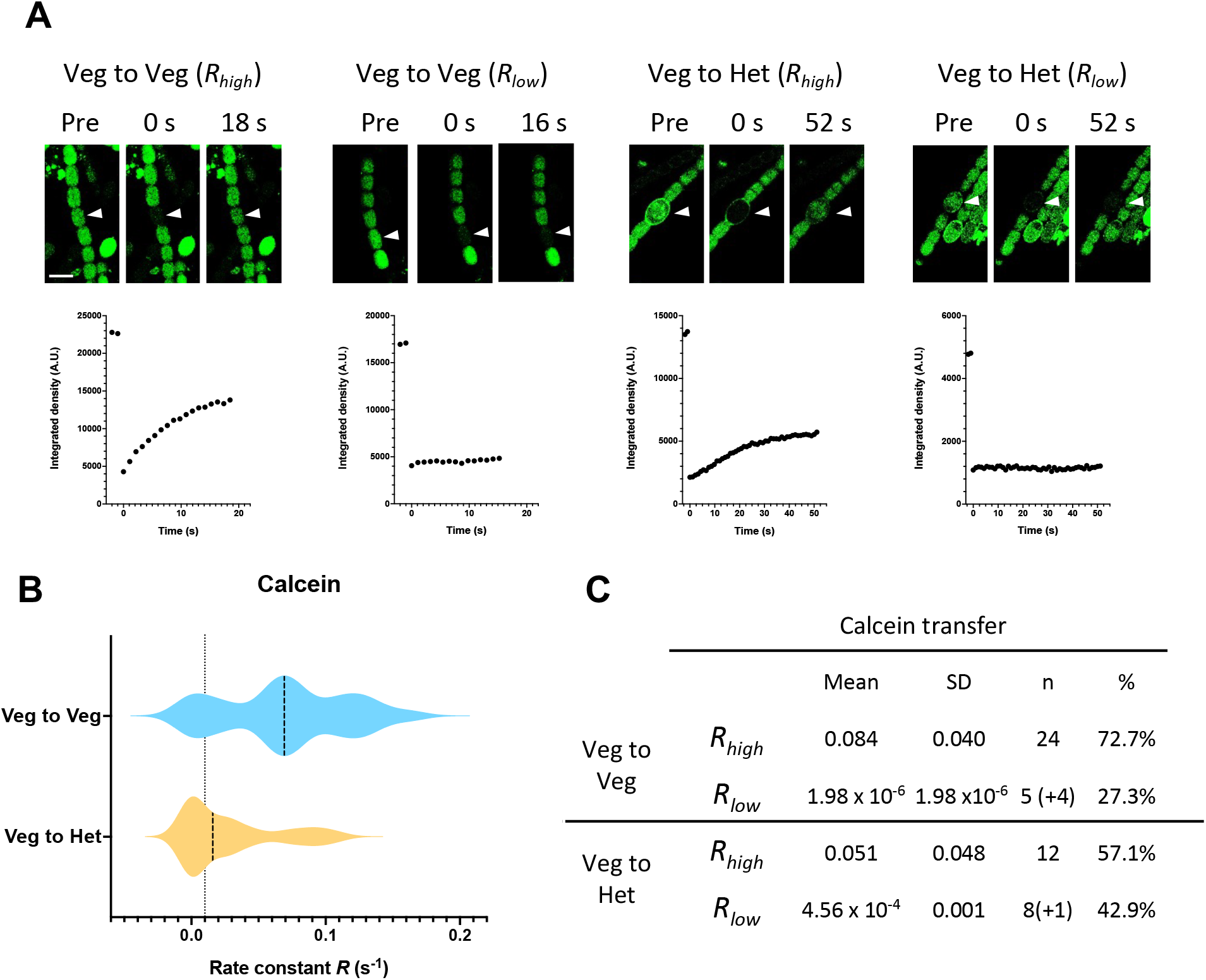
Calcein FRAP analysis of *N. azollae* filaments. (A) Calcein FRAP experiments. Filaments were labelled and subjected to FRAP analysis as described in Materials and Methods. The cells indicated by arrowheads were photobleached and their fluorescence was monitored continuously. Example of images prior (pre), and at 0 s and 16-52 s after bleaching are shown in the upper panel. Scale bars, 10 μm. Non-normalized fluorescence recovery curves for their respective bleached cells are shown below. (B) Violin Plot of recovery rate constants (*R*) for calcein recovery between vegetative cells (Veg to Veg) and from vegetative cells to a heterocyst (Veg to Het). The dotted line indicates the threshold, *R =* 0.01 s^−1^, that is used to discriminate two population of cells, communicating cells (*R*_*high*_, *R* ≥ 0.01 s^−1^) and noncommunicating cells (*R*_*low*_, *R <* 0.01 s^−1^). Dashed lines depict the median values. (C) Table summarizing the mean rate constants grouped by communicating (*R*_*high*_) and noncommunicating (*R*_*low*_) cells observed for calcein transfer. Numbers in parentheses represent noncommunicating cells that did not adjust to the model making it not possible to obtain an *R* number, but are included to calculate a percentage.

The transfer of calcein between vegetative cells, took place in about 73 % of the cells tested with a recovery constant, *R*, of about 0.084 s^−1^ (Fig. 2B, 2C). This is similar to values described for free-living *Anabaena* grown in the presence of nitrate or incubated in the absence of a source of combined nitrogen (17, 25, 33, 38). The rest of the cells examined (27 %) showed *R* < 0.01 s^−1^, which indicates that they were noncommunicating cells as defined previously (33). Transfer of calcein from a vegetative cell to a heterocyst was found to take place in about 57 % of the heterocysts with an *R* value of about 0.051 s^−1^, which is lower than between vegetative cells as has been described for *Anabaena* (17). The rest of the heterocysts tested were noncommunicating (43 %, *R* < 0.01 s^−1^).

We also tested 5-CF transfer between vegetative cells and from vegetative cells to heterocysts. About 84 % of the vegetative cells tested were communicating with a mean *R* value of 0.08 s^−1^, which is identical to a value described for *Anabaena* (25), with about 16 % of the vegetative cells being noncommunicating (Fig. 3B, 3C). 5-CF transfer to heterocysts took place only in about 27 % of the heterocysts tested, and the observed *R* value was relatively low, 0.040 s^−1^. Transfer of 5-CF from vegetative cells to heterocysts has not been reported before for any heterocyst-forming cyanobacterium, but our results show that this marker can also be used to test molecular transfer between the different cell types (vegetative cells and heterocysts).

**Figure 3.**
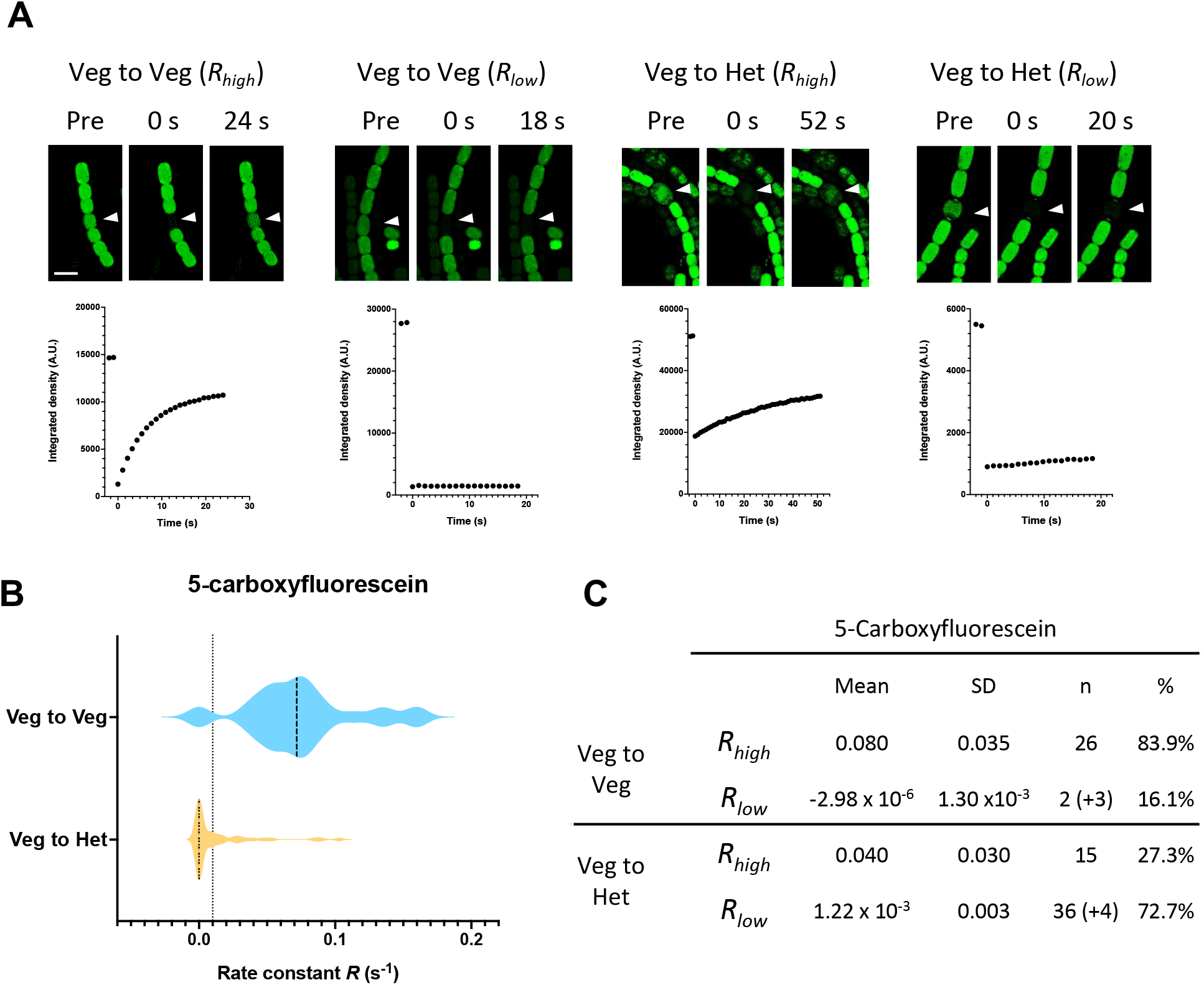
5-Carboxyfluorescein FRAP analysis of *N. azollae* filaments. (A) 5-CF FRAP experiments. Filaments were labelled with 5-CF and subjected to FRAP analysis as described in Materials and Methods. The cells indicated by arrowheads were photobleached and their fluorescence was monitored continuously. Example of images prior (pre), and at 0 s and 16-52 s after bleaching are shown in the upper panel. Scale bars, 10 μm. Non-normalized fluorescence recovery curves for their respective bleached cells are shown below. (B) Violin Plot of recovery rate constants (*R*) for 5-CF recovery between vegetative cells (Veg to Veg) and from vegetative cells to a heterocyst (Veg to Het). The dotted line indicates the threshold, *R =* 0.01 s^−1^, that is used to discriminate two population of cells, communicating cells (*R*_*high*_, *R* ≥ 0.01 s^−1^) and noncommunicating cells (*R*_*low*_, *R <* 0.01 s^−1^). Dashed lines depict the median values. (C) Table depicting the mean rate constants grouped by communicating cells (*R*_*high*_) and noncommunicating cells (*R*_*low*_) observed for 5-CF transfer. Numbers in parentheses represent the noncommunicating cells that did not adjust to the model making it not possible to obtain an *R* number, but are included to calculate a percentage.

The high percentage of noncommunicating heterocysts for 5-CF was also noticed by the observation of non-stained heterocysts before the FRAP routine, as in several cases a heterocyst within a filament exhibited weak or no fluorescence when stained with 5-CF, in spite of being surrounded by labeled vegetative cells (Suppl. Fig. S2). This result indicates a low or null permeability of the heterocyst cell wall to 5-CF and suggests that the heterocysts labeled received 5-CF from their adjacent vegetative cells. Because, as shown with esculin (25), heterocysts lose communication as they senesce, the relatively low number of communicating heterocysts detected with 5-CF likely reflects the heterogeneity of *N. azollae* cells in the *Azolla* juice. The percentage of communicating heterocysts detected with calcein was however larger than that determined with 5-CF (Figs. 2C and 3C), consistent with the idea that calcein and 5-CF transfer are not completely equivalent and different types of septal junctions may have preference for one or the other marker (25, 39, 40).

### *Nostoc azollae* contains a rich array of septal nanopores

Murein (PG) sacculi were isolated from filaments/cells in the *Azolla* juice and prepared and visualized by transmission electron microscopy (TEM) as described in Materials and Methods. Murein sacculi corresponding to one or more cells were observed that contained septal disks (Fig. 4A). The septal PG disks had a mean diameter of about 1.6 μm, which is larger than that described for *Anabaena* (about 0.9 μm; (22)), but disks clearly smaller and larger than that mean size could be observed (Fig. 4B).

**Figure 4.**
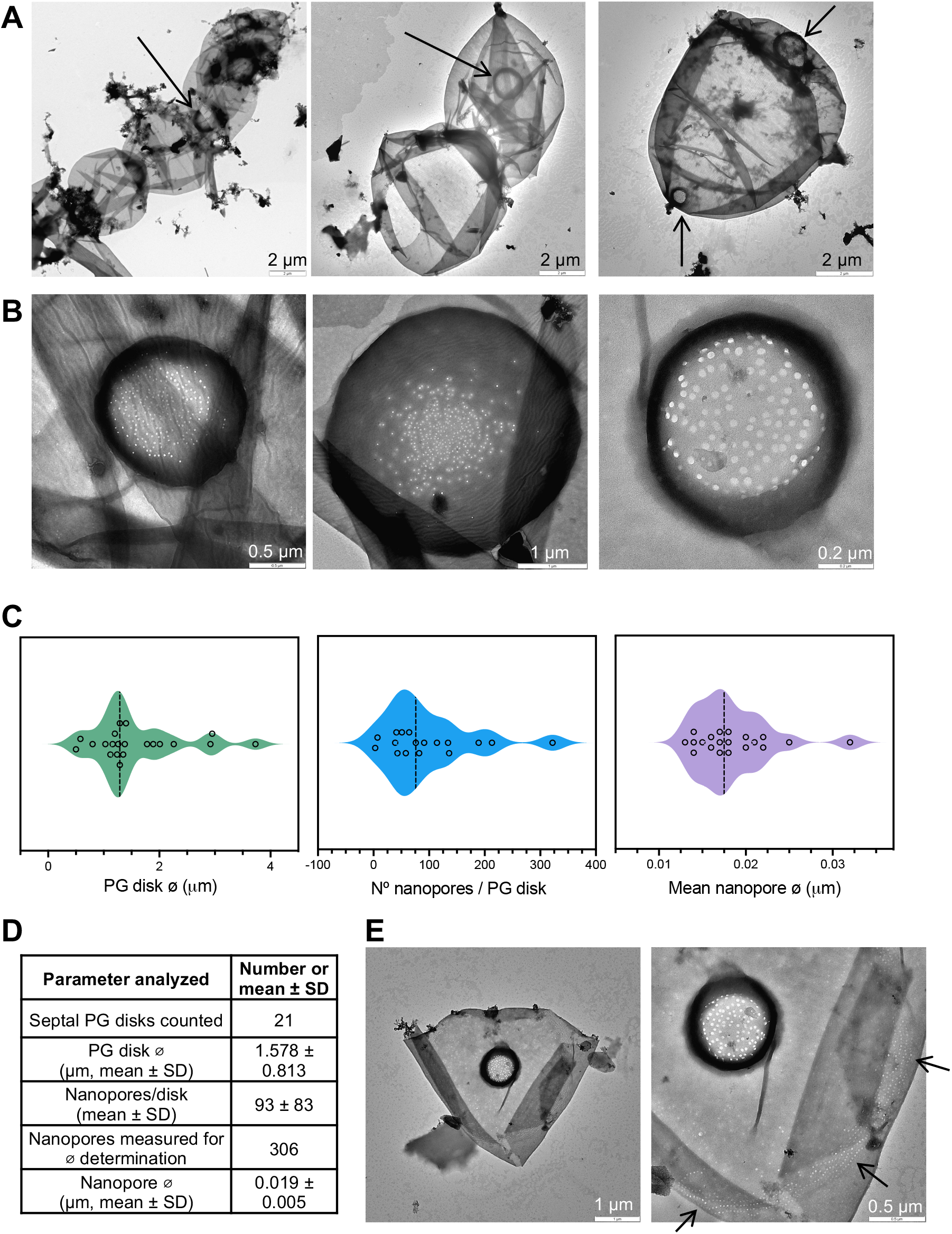
Nanopores in septal peptidoglycan disks of *N. azollae* filaments. (A) Murein sacculi of a fragment of a filament (*left*), of two cells (*middle*) and of a single cell (separated from other cells in the filament; *right*); the arrows point to septal PG disks. (B) Examples of septal disks and corresponding nanopore arrays for an average diameter disk, ⌀ 1.38 μm (*left*), an over-average disk, ⌀ 3.73 μm (*middle*), and a lower-than average disk, ⌀ 0.80 μm (*right*). The PG was isolated and visualized by TEM as described in Materials and Methods. Scale bars are shown. (C) Violin plots depicting the measurements of PG disk diameter (μm), number of nanopores/disk, and nanopore diameters. Dashed lines depict the median values. (D) Table summarizing the means and standard deviations of nanopore numbers per septal PG disk and diameters of PG disks and nanopores. (E) The sacculus shown at different magnifications may correspond to a heterocyst because it contains “arcs of small pores” (arrows).

All the disks observed contained nanopores, with 93 ± 83 nanopores per septal disk (mean ± SD), indicating a wide variability (Fig. 4D). In other studies, 100 to 250 nanopores (also termed “microplasmodesmata”) per septal disk were reported for *Anabaena variabilis* and *Anabaena cylindrica* (41), about 155 for *Nostoc punctiforme* (23), and about 75 (25) or 41 (33) for *Anabaena*. The mean diameter of the *N. azollae* nanopores was about 19 nm (Fig. 4C), similar to *Anabaena* (25). Because there was some variability in nanopore number and disk and nanopore diameters, we examined possible correlations between these parameters. There was not a significant correlation between nanopore number and nanopore diameter (R^2^ = 0.124; *P* = 0.165) nor between septal disk and nanopore size (R^2^ = 0.109; *P* = 0.160), whereas a moderate positive correlation was observed between nanopore number and septal disk diameter (R^2^ = 0.277; *P* = 0.030) (Suppl. Fig. S3).

In two cases, arcs of small pores were observed in the sacculus outside of the septal disks (one example shown in Fig. 4E). These arcs of small pores have been described in heterocyst sacculi of *Anabaena*, where they have been suggested to be involved in polysaccharide translocation across the PG (22). These results are in good agreement with the presence of orthologues of key *Anabaena* proteins involved in intercellular molecular transfer in the *N. azollae* genome. Specifically with the conservation of amidases AmiC1, AmiC2, and a LytM factor in *N. azollae*, which are important for the formation of nanopores in septal disks (42, 43) (Table 1).

### Concluding remarks

The results presented here show that the juice prepared from *Azolla* plants contains live *Nostoc azollae* cells, as deduced from the observation that *N. azollae* cells freshly isolated from the fern are metabolically active. This is consistent with previous work in which freshly isolated *N. azollae* cells were used to study, e.g., [^13^N]N_2_ fixation and the fate of fixed ^13^N (15). Most of the extracted *N. azollae* was largely present as filaments, and many filaments contained heterocysts. FRAP analysis showed substantial activities of intercellular transfer of calcein and 5-CF, indicating that the cells in the *N. azollae* filaments can communicate with each other. This aligns with the presence in the eroded genome of *N. azollae* of orthologues for a majority of the key proteins involved in the functionality of the intercellular molecular transfer machinery. Intercellular communication is essential for growth of the filaments, in which the heterocysts provide the vegetative cells with fixed N and the vegetative cells provide heterocysts with reduced C. Additionally, noncommunicating vegetative cells were observed, indicating that the mechanism(s) that regulate septal junction opening/closing in *Anabaena* (20), with closing making a cell noncommunicating (33), are conserved in *N. azollae*. Noncommunicating heterocysts were also observed, which may correspond to senescent heterocysts as previously discussed (25). Thus, in spite of an endophytic lifestyle, *N. azollae* filaments retain a fully active intercellular communication system that allows its growth as a multicellular organism. On the other hand, the range of sizes of the septal PG disks is wider in *N. azollae* than in the *Anabaena* model (22). This observation may reflect the presence in the *Azolla* juice of different cell types, including the vegetative cells and heterocysts of diazotrophic filaments and the cells of hormogonia. The hormogonial cells are smaller than those in diazotrophic filaments, likely involving smaller intercellular septa and, thus, smaller PG disks. Additionally, in diazotrophic filaments, vegetative cell-vegetative cell septa and vegetative cell-heterocyst septa may also have different sized PG disks, which may contribute to the different rates of fluorescent marker transfer observed from vegetative cells to heterocysts and between vegetative cells.

## Materials and Methods

### *Azolla* fern culturing

This study used *Azolla filiculoides*, strain Galgenwaard (2). Adult *Azolla* sporophytes were grown in a modified SAM medium, a variation of IRRI2 medium (44), under a 16-h light-8 h dark diel cycle at 20 °C. To maintain optimal growth conditions, white light (100 μmol m^−2^ s^−1^) was supplemented with Far-Red light from an *APEXstrip Bundle 730 nm −16W system (Crescience)*.

### *Azolla* juice extraction

*N. azollae*-enriched *Azolla* juice was obtained following the protocol visually documented in Supplementary Figure S1. Briefly, an amount of ~10 g of *Azolla* fronds was placed in dH_2_O-rinsed ziplock bags and mixed with 10 mL of BG11_0_ medium. The juice was extracted by squeezing the ziplock bags containing the fronds using a pasta maker machine by gently crush the plant to minimize the release of oxidants. The juice obtained from 2-3 extractions was gently centrifuged at 3000 × *g* for 5 min at room temperature. The supernatant was discarded and the resulting pellet, containing material enriched in *N. azollae* was resuspended in half of the total extraction volume of fresh BG11_0_ medium (45). For nanopore analysis, pelleted cells were extracted in dH_2_O instead of BG11_0_ medium and the pellet was stored at −20 °C until PG isolation.

### Labelling with the sucrose-analogue esculin

Esculin labeling was performed with a protocol adapted from the previously described by (25) cells from a 1-mL aliquot of the *N. azollae*-enriched *Azolla* juice suspension (obtained as described above) were harvested by gentle centrifugation at room temperature, resuspended in 500 μL of fresh BG11_0_ medium, and mixed with 15 μL of a saturated esculin hydrate solution (~5 mM, Sigma-Aldrich) in water. The mixture was then incubated in the dark for 1 h at 30 °C with gentle shaking, followed by three washes with BG11_0_ medium. The cells were subsequently incubated in the dark for another 15 min in 1 mL of medium at 30 °C with mild agitation. After this, the cells were washed again, concentrated about 20-fold, and applied (20 to 40 μL) onto BG11_0_ medium plates solidified with 1% (wt/vol) Difco Bacto agar. Pieces of agar from the BG11_0_ medium plate carrying cells/filaments were cut out and covered with a coverslip for visualization by confocal microscopy. Esculin was excited at 355 nm and fluorescence was detected within the 443 to 490 nm range.

### FRAP experiments

Calcein and 5-Carboxyfluorescein (5-CF) labeling was performed as previously reported by Mullineaux *et al*. (17) and Merino-Puerto *et al*. (39), respectively. Briefly, for calcein staining, cells from 0.5 mL of *N. azollae*-enriched *Azolla* juice suspension (obtained as described above extracted in BG11_0_ medium) were harvested by gentle centrifugation, washed two times, resuspended in 0.5 mL of fresh BG11_0_ medium, and mixed with 10 μL of calcein-AM (1 mg/mL in dimethyl sulfoxide). The suspensions were incubated in the dark at 30 °C for 90 min, and cells were then harvested and washed three times in fresh, dye-free BG11_0_ medium and resuspended in 175 μL of dye-free BG11_0_ medium. For 5-CF, 1 mL of *N. azollae*-enriched *Azolla* juice suspension was harvested, washed two times and resuspended in 1 mL of fresh BG11_0_ medium, and then mixed with 4 μL of 5-carboxyfluorescein diacetate AM (5 mg/mL in dimethyl sulfoxide). The suspension was incubated in the dark at 30 °C for 30 min, and filaments were then harvested and washed three times in fresh, dye-free BG11_0_ medium and resuspended in 250 μL of dye-free BG11_0_ medium.

Cell suspensions from calcein and 5-CF labelling were spotted and spread onto 30 mL-plates of BG11_0_ medium agar and placed in a temperature-controlled sample holder with a glass cover slip on top. All measurements were carried out at 30 °C. Images were collected with an Olympus FLUOVIEW FV3000 confocal laser-scanning microscope equipped with a UPlanApo 60 × 1.5 NA oil immersion objective. Excitation was achieved with a 488-nm argon laser and fluorescent emission was monitored by collection across windows of 500 to 520 nm. After an initial image was recorded, the bleach was carried out by a pre-set FRAP routine previously described (17) using 20% laser and occasionally 40% laser for cases when total bleach was difficult to obtain (large-sized heterocysts). In all cases, values of fluorescence recovery were normalized to the prebleached value. Kinetics of transfer of calcein and 5-CF was computed with the Fiji processing package from ImageJ (46) and the recovery rate constant, *R*, was calculated as previously described (26) for both fluorescent markers.

### Nanopore analysis

Cells from *N. azollae*-enriched *Azolla* juice suspensions were obtained as described above and PG sacculi were isolated and analyzed as previously described (23, 25). The purified PG sacculi were placed on formvar/carbon film coated 150-mesh copper grids (*Electron Microscopy Sciences*) and stained with 1% (w/v) uranyl acetate. Samples were visualized using a Zeiss Libra 120 Plus electron microscope at 120 kV.

## Acknowledgments

Work was supported by the Gordon and Betty Moore Foundation grant no. 9355.

## Author contributions

CSB designed and performed experimental work and interpreted data; MNM provided methodology; PL, SNB, HS raised funds and supervised work; EF supervised work and drafted the manuscript; all authors provided input on the content of the manuscript and the figures.

## Notes

### Competing Interest Statement

The authors have declared no competing interest.

## References

1. Brouwer P, Bräutigam A, Buijs VA, Tazelaar AOE, van der Werf A, Schlüter U, Reichart GJ, Bolger A, Usadel B, Weber APM, Schluepmann H. 2017. Metabolic adaptation, a specialized leaf organ structure and vascular responses to diurnal N_2_ fixation by nostoc azollae sustain the astonishing productivity of Azolla ferns without nitrogen fertilizer. Front Plant Sci 8.

2. Li FW, Brouwer P, Carretero-Paulet L, Cheng S, De Vries J, Delaux PM, Eily A, Koppers N, Kuo LY, Li Z, Simenc M, Small I, Wafula E, Angarita S, Barker MS, Bräutigam A, Depamphilis C, Gould S, Hosmani PS, Huang YM, Huettel B, Kato Y, Liu X, Maere S, McDowell R, Mueller LA, Nierop KGJ, Rensing SA, Robison T, Rothfels CJ, Sigel EM, Song Y, Timilsena PR, Van De Peer Y, Wang H, Wilhelmsson PKI, Wolf PG, Xu X, Der JP, Schluepmann H, Wong GKS, Pryer KM. 2018. Fern genomes elucidate land plant evolution and cyanobacterial symbioses. Nat Plants 4:460.

3. Peters GA, Mayne BC, Kettering CF. 1974. The Azolla, Anabaena azollae Relationship: I. Initial Characterization of the Association. Plant Physiol 53:813–819.

4. Dijkhuizen LW, Brouwer P, Bolhuis H, Reichart GJ, Koppers N, Huettel B, Bolger AM, Li FW, Cheng S, Liu X, Wong GKS, Pryer K, Weber A, Bräutigam A, Schluepmann H. 2018. Is there foul play in the leaf pocket? The metagenome of floating fern Azolla reveals endophytes that do not fix N_2_ but may denitrify. New Phytologist 217:453–466.

5. Ran L, Larsson J, Vigil-Stenman T, Nylander JAA, Ininbergs K, Zheng WW, Lapidus A, Lowry S, Haselkorn R, Bergman B. 2010. Genome erosion in a nitrogen-fixing vertically transmitted endosymbiotic multicellular cyanobacterium. PLoS One 5.

6. Zheng W, Bergman B, Chen B, Zheng S, Xiang G, Rasmussen U. 2009. Cellular responses in the cyanobacterial symbiont during its vertical transfer between plant generations in the Azolla microphylla symbiosis. New Phytologist 181:53–61.

7. Hill DJ. 1975. The pattern of development of Anabaena in the Azolla-Anabaena symbiosis. Planta 122:179–184.

8. Canini A, Grilli Caiola M, Mascini M. 1990. Ammonium content, nitrogenase activity and heterocyst frequency within the leaf cavities of Azolla filiculoides Lam. FEMS Microbiol Lett 71:205–210.

9. Zheng W, Bergman B, Chen B, Zheng S, Xiang G, Rasmussen U. 2009. Cellular responses in the cyanobacterial symbiont during its vertical transfer between plant generations in the Azolla microphylla symbiosis. New Phytologist 181:53–61.

10. Kaplan D, Calvert HE, Peters GA. 1986. The Azolla-Anabaena azollae Relationship : XII. Nitrogenase Activity and Phycobiliproteins of the Endophyte as a Function of Leaf Age and Cell Type. Plant Physiol 80:884– 890.

11. Kaplan D, Peters GA. 1981. The Azolla-Anabaena azollae relationship. New Phytologist 89:337–346.

12. Hill DJ. 1977. The role of Anabaena in the Azolla-Anabaena symbiosis. New Phytologist 78:611–616.

13. Braun-Howland EB, Lindblad P, Nierzwicki-Bauer SA, Bergman B. 1988. Dinitrogenase reductase (Fe-protein) of nitrogenase in the cyanobacterial symbionts of three Azolla species: Localization and sequence of appearance during heterocyst differentiation. Planta 176:319–322.

14. Nierzwicki-Bauer SA, Haselkorn R. 1986. Differences in mRNA levels in Anabaena living freely or in symbiotic association with Azolla. EMBO J 5:29–35.

15. Meeks JC, Steinberg N, Joseph CM, Enderlin CS, Jorgensen PA, Peters GA. 1985. Assimilation of exogenous and dinitrogen derived 13NH4_+_ by Anabaena azollae separated from Azolla caroliniana Willd. Archives of microbiology, Springer 142:229–233.

16. Nieves-Morión M, Flores E, Whitehouse MJ, Thomen A, Foster RA. 2021. Single-cell measurements of fixation and intercellular exchange of C and N in the filaments of the heterocyst-forming cyanobacterium Anabaena sp. Strain PCC 7120. mBio 12.

17. Mullineaux CW, Mariscal V, Nenninger A, Khanum H, Herrero A, Flores E, Adams DG. 2008. Mechanism of intercellular molecular exchange in heterocyst-forming cyanobacteria. EMBO J 27:1299.

18. Nieves-Morión M, Mullineaux CW, Flores E. 2017. Molecular Diffusion through Cyanobacterial Septal Junctions. mBio 8:e01756–16.

19. Flores E, Nieves-Morión M, Mullineaux CW. 2018. Cyanobacterial Septal Junctions: Properties and Regulation. Life (Basel) 9.

20. Weiss GL, Kieninger AK, Maldener I, Forchhammer K, Pilhofer M. 2019. Structure and Function of a Bacterial Gap Junction Analog. Cell 178:374-384.e15.

21. Wilk L, Strauss M, Rudolf M, Nicolaisen K, Flores E, Kühlbrandt W, Schleiff E. 2011. Outer membrane continuity and septosome formation between vegetative cells in the filaments of Anabaena sp. PCC 7120. Cell Microbiol 13:1744–1754.

22. Arévalo S, Flores E. 2021. Heterocyst Septa Contain Large Nanopores That Are Influenced by the Fra Proteins in the Filamentous Cyanobacterium Anabaena sp. Strain PCC 7120. J Bacteriol 203.

23. Lehner J, Berendt S, Dörsam B, Pérez R, Forchhammer K, Maldener I. 2013. Prokaryotic multicellularity: a nanopore array for bacterial cell communication. FASEB J 27:2293–2300.

24. Nierzwicki-Bauer SA, Aulfinger H. 1991. Occurrence and ultrastructural characterization of bacteria in association with and isolated from Azolla caroliniana. Appl Environ Microbiol 57:3629–3636.

25. Nürnberg DJ, Mariscal V, Bornikoel J, Nieves-Morión M, Krauß N, Herrero A, Maldener I, Flores E, Mullineaux CW. 2015. Intercellular Diffusion of a Fluorescent Sucrose Analog via the Septal Junctions in a Filamentous Cyanobacterium 10.1128/mBio.02109-14.

26. Nieves-Morión M, Lechno-Yossef S, López-Igual R, Frías JE, Mariscal V, Nürnberg DJ, Mullineaux CW, Wolk CP, Flores E. 2017. Specific glucoside transporters influence septal structure and function in the filamentous, heterocyst-forming cyanobacterium Anabaena sp. strain PCC 7120. J Bacteriol 199:876–892.

27. Flores E, Nieves-Morión M, Mullineaux CW. 2018. Cyanobacterial Septal Junctions: Properties and Regulation. Life (Basel) 9.

28. Nieves-Morión M, Flores E. 2018. Multiple ABC glucoside transporters mediate sugar-stimulated growth in the heterocyst-forming cyanobacterium Anabaena sp. strain PCC 7120. Environ Microbiol Rep 10:40–48.

29. López-Igual R, Flores E, Herrero A. 2010. Inactivation of a heterocyst-specific invertase indicates a principal role of sucrose catabolism in heterocysts of Anabaena sp. J Bacteriol 192:5526–5533.

30. Vargas WA, Nishi CN, Giarrocco LE, Salerno GL. 2011. Differential roles of alkaline/neutral invertases in Nostoc sp. PCC 7120: Inv-B isoform is essential for diazotrophic growth. Planta 233:153–162.

31. Kaplan D, Peters GA, Kaplan D, Peters GA. 1988. Interaction of Carbon Metabolism in the Azolla-Anabaena Symbiosis. Symbiosis. Balaban Publishers.

32. Flores E, Pernil R, Muro-Pastor AM, Mariscal V, Maldener I, Lechno-Yossef S, Fan Q, Wolk CP, Herrero A. 2007. Septum-localized protein required for filament integrity and diazotrophy in the heterocyst-forming cyanobacterium Anabaena sp. strain PCC 7120. J Bacteriol 189:3884– 3890.

33. Arévalo S, Nenninger A, Nieves-Morión M, Herrero A, Mullineaux CW, Flores E. 2021. Coexistence of Communicating and Noncommunicating Cells in the Filamentous Cyanobacterium Anabaena. mSphere 6.

34. Kieninger AK, Tokarz P, Janović A, Pilhofer M, Weiss GL, Maldener I. 2022. SepN is a septal junction component required for gated cell–cell communication in the filamentous cyanobacterium Nostoc. Nature Communications 2022 13:1 13:1–15.

35. Rudolf M, Tetik N, Ramos-León F, Flinner N, Ngo G, Stevanovic M, Burnat M, Pernil R, Flores E, Schleiffa E. 2015. The Peptidoglycan-Binding Protein SjcF1 Influences Septal Junction Function and Channel Formation in the Filamentous Cyanobacterium Anabaena. mBio 6:e00376–15.

36. Schätzle H, Arévalo S, Flores E, Schleiff E. 2021. A TonB-Like Protein, SjdR, Is Involved in the Structural Definition of the Intercellular Septa in the Heterocyst-Forming Cyanobacterium Anabaena. mBio 12.

37. Kamiloglu S, Sari G, Ozdal T, Capanoglu E. 2020. Guidelines for cell viability assays. Food Front 1:332–349.

38. Mariscal V, Herrero A, Nenninger A, Mullineaux CW, Flores E. 2011. Functional dissection of the three-domain SepJ protein joining the cells in cyanobacterial trichomes. Mol Microbiol 79:1077–1088.

39. Merino-Puerto V, Schwarz H, Maldener I, Mariscal V, Mullineaux CW, Herrero A, Flores E. 2011. FraC/FraD-dependent intercellular molecular exchange in the filaments of a heterocyst-forming cyanobacterium, Anabaena sp. Mol Microbiol 82:87–98.

40. Mariscal V, Nürnberg DJ, Herrero A, Mullineaux CW, Flores E. 2016. Overexpression of SepJ alters septal morphology and heterocyst pattern regulated by diffusible signals in Anabaena. Mol Microbiol 10.1111/mmi.13436.

41. Giddings TH, Staehelin LA. 1981. Observation of microplasmodesmata in both heterocyst-forming and non-heterocyst forming filamentous cyanobacteria by freeze-fracture electron microscopy. Archives of Microbiology 1981 129:4 129:295–298.

42. Bornikoel J, Staiger J, Madlung J, Forchhammer K, Maldener I. 2018. LytM factor Alr3353 affects filament morphology and cell–cell communication in the multicellular cyanobacterium Anabaena sp. PCC 7120. Mol Microbiol 108:187–203.

43. Bornikoel J, Carrión A, Fan Q, Flores E, Forchhammer K, Mariscal V, Mullineaux CW, Perez R, Silber N, Wolk CP, Maldener I. 2017. Role of Two Cell Wall Amidases in Septal Junction and Nanopore Formation in the Multicellular Cyanobacterium Anabaena sp. 7:1–15.

44. Dijkhuizen LW, Tabatabaei BES, Brouwer P, Rijken N, Buijs VA, Güngör E, Schluepmann H. 2021. Far-Red Light-Induced Azolla filiculoides Symbiosis Sexual Reproduction: Responsive Transcripts of Symbiont Nostoc azollae Encode Transporters Whilst Those of the Fern Relate to the Angiosperm Floral Transition. Front Plant Sci 12:693039.

45. Rippka R, Deruelles J, Waterbury JB. 1979. Generic assignments, strain histories and properties of pure cultures of cyanobacteria. J Gen Microbiol 111:1–61.

46. Schindelin J, Arganda-Carreras I, Frise E, Kaynig V, Longair M, Pietzsch T, Preibisch S, Rueden C, Saalfeld S, Schmid B, Tinevez JY, White DJ, Hartenstein V, Eliceiri K, Tomancak P, Cardona A. 2012. Fiji: an open-source platform for biological-image analysis. Nature Methods 2012 9:7 9:676–682.

47. Janović A, Maldener I, Menzel C, Parrett GA, Risser DD. 2024. The role of FraI in cell-cell communication and differentiation in the hormogonia-forming cyanobacterium Nostoc punctiforme. mSphere 9.

48. Arévalo S, Flores E. 2020. Pentapeptide-repeat, cytoplasmic-membrane protein HglK influences the septal junctions in the heterocystous cyanobacterium Anabaena. Mol Microbiol 113:794–806.

49. Springstein BL, Arévalo S, Helbig AO, Herrero A, Stucken K, Flores E, Dagan T. 2020. A novel septal protein of multicellular heterocystous cyanobacteria is associated with the divisome. Mol Microbiol 113:1140– 1154.

50. Velázquez-Suárez C, Springstein BL, Nieves-Morión M, Helbig AO, Kieninger AK, Maldener I, Nürnberg DJ, Stucken K, Luque I, Dagan T, Herrero A. 2023. SepT, a novel protein specific to multicellular cyanobacteria, influences peptidoglycan growth and septal nanopore formation in Anabaena sp. PCC 7120. mBio 14.

51. Sarasa-Buisan C, Nieves-Morión M, Arévalo S, Helm RF, Sevilla E, Luque I, Fillat MF. 2024. FurC (PerR) contributes to the regulation of peptidoglycan remodeling and intercellular molecular transfer in the cyanobacterium Anabaena sp. strain PCC 7120. mBio 15.

