## Supplementary material for "Intercellular communication in the fern endosymbiotic cyanobacterium *Nostoc azollae*"

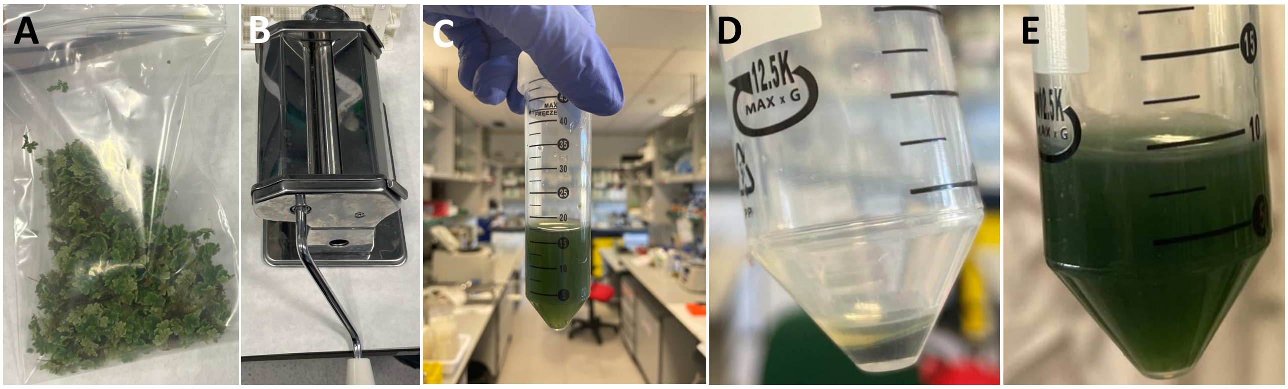


**Figure S1. Extraction of *Nostoc azollae*-enriched "*Azolla* juice" from *Azolla filiculoides*, strain Galgenward.** A, 10 g FW *Azolla* filliculoides strain Galgenward was transferred in a zip=lock plastic bag. B, Pastamaker device to gently squeeze the *Azolla* fronds inside the zip-lock bag. C, "*Azolla* juice" obtained after collection of the material in BG11$_0$ medium. D, Pellet enriched in *N. azollae* cells. E, Final *N. azollae* enriched suspension in BG11$_0$ medium for esculin labelling and FRAP-analysis.

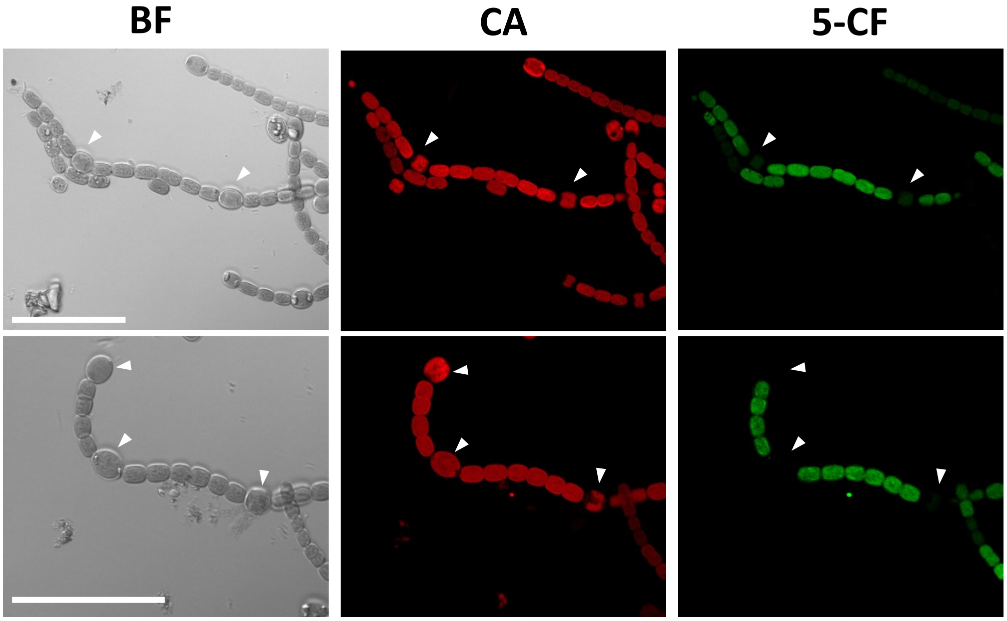


**Figure S2. Absence of 5-CF fluorescence in some heterocysts before photobleaching during the FRAP routine.** Filaments were labelled with 5-CF (5-Carboxyfluorescein) as described in Materials and Methods. The heterocysts indicated by arrowheads were not photobleached. Scale bars, 50 μm. The two rows show different examples, in bright field (BF), and fluorescence emission for cyanobacterial autofluorescence (CA) and 5-CF fluorescence (5-CF).

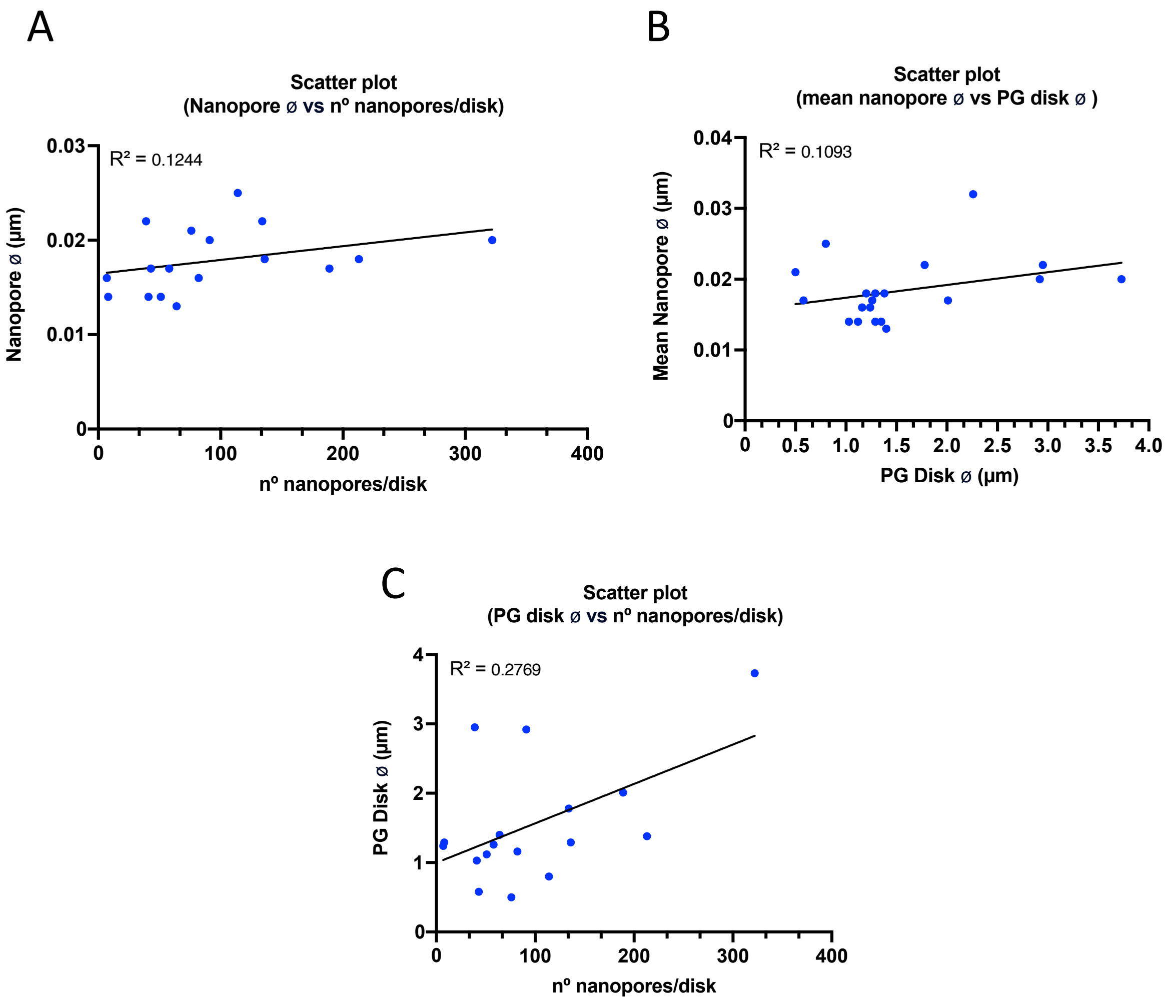


**Figure S3.** Scatter plots showing (A) nanopore diameters vs. number of nanopores/disk, (B) nanopore vs. PG-disk diameters, and (C) PG-disk diameters vs. number of nanopores/disk. Pearson's correlation coefficient ($R^2$) is shown in each case.
